# Synaptic transmission may provide an evolutionary benefit to HIV through modulation of latency

**DOI:** 10.1101/243360

**Authors:** Cesar Vargas-Garcia, Ryan Zurakowski, Abhyudai Singh

## Abstract

Transmission of HIV is known to occur by two mechanisms in vivo: the free virus pathway, where viral particles bud off an infected cell before attaching to an uninfected cell, and the cell-cell pathway, where infected cells form virological synapses through close contact with an uninfected cell. It has also been shown that HIV replication includes a positive feedback loop controlled by the viral protein Tat, which may act as a stochastic switch in determining whether an infected cell enters latency. In this paper, we introduce a simple mathematical model of HIV replication containing both the free virus and cell-cell pathways. Using this model, we demonstrate that the high multiplicity of infection in cell-cell transmission results in a suppression of latent infection, and that this modulation of latency through balancing the two transmission mechanisms can provide an evolutionary benefit to the virus. This benefit increases with decreasing overall viral fitness, which may provide a within-host evolutionary pressure toward more cell-cell transmission in late-stage HIV infection.

## Introduction

The Human Immunodeficiency Virus (HIV) is a retrovirus that primarily targets the CD4+ T-cells. While it is susceptible to antiviral compounds at several points during its life cycle, it is capable of creating replication-competent chromosome-integrated infections with very little viral transcription [7, 6]. These so-called quiescently-infected cells are not affected by any of the current antiviral compounds [4]. Such quiescent or latent cells can persist for decades before randomly transitioning to a fully activated, virus producing state, and constitutes a major barrier to eradicating the virus from a patient [23, 28, 24].

The HIV cell-fate decision of whether to follow a latent or active-infection pathway is critically controlled through a positive-feedback loop mediated by the Tat viral protein [30, 34, 21, 29, 10]. Tat facilitates the successful transcription of the integrated HIV genome [15]. When Tat is suppressed, little to no viral transcription is observed [18]. Conversely, when exogenous Tat is supplied, latency is suppressed [5]. Once successful transcription has occurred, translation provides an adequate number of Tat molecules to facilitate continued transcription and translation, providing the positive feedback mechanism.

As HIV transcription, translation, and assembly progress in an infected cell, the HIV surface molecules gp120 and gp41 begin to accumulate on the surface of the infected cell [25]. These molecules have a high affinity for the CD4 and CCR5 molecules on T Cells, and facilitate the binding and membrane fusion processes during infection by free virions. However, the molecules on the surface of the infected cell can also facilitate the formation of synapses between infected cells and uninfected T Cells [19]. This is recognized as a major secondary pathway for HIV transmission, in addition to the free virus pathway [27].

The synaptic pathway results in the equivalent of a large number of virions being deposited into the target cell; this is referred to as multiplicity of infection. While this increases the probability of a successful infection, it does so at the cost of forgoing the possibility of infecting other target cells via the free virus pathway. Potential evolutionary advantages and disadvantages of utilizing the synaptic pathway of transmission have been previously examined in several works [12, 27, 13, 11, 14, 33]. We propose a novel explanation for the evolution of the synaptic pathway; the modulation of the probability of latency.

Modulation of latency through the balance of synaptic and free virus transmissions pathways occurs in the following way. An initial, randomly distributed number of Tat molecules are transmitted during infection. These molecules may reside in the virion [1], or they may be secreted by the parent cell and subsequently endogenized by the target cell [3]. The more Tat molecules are present, the more likely that transcription will complete before the initial Tat molecule population degrades. Cells infected via the synaptic pathway have many times more Tat molecules present at infection than cells infected via the free virus pathway. This dramatically reduces the probability that cells infected by the synaptic pathway will enter a latent state.

In this paper, we introduce a simple mathematical model of HIV replication that accounts for both the free virus and cell-cell transmission pathways. The dynamics include the effect described above, namely that the likelihood of an infected cell entering a latent state is reduced if that cell was infected via the cell-cell pathway as compared to a cell infected by the free virus pathway. We demonstrate through mathematical modeling that viruses with the cell-cell transmission pathway have a selective advantage compared to viruses without this mechanism, with an optimal fraction of total virus transmitted through the synaptic pathway ranging between 0% and 20%, depending primarily on the per virus probability of infection, the probability that a cell infected by a single virus will enter latency, and the basic reproductive ratio of the free-virus pathway. We also show that the optimal fraction of viruses sent through the synaptic pathway dramatically increases as the basic reproductive ratio of the free virus pathway decreases. We hypothesize that this may create a pressure for within-host evolution toward virus that promotes cell-cell binding, which may lead to syncytia formation in late-stage HIV disease.

## HIV Model

The free virus transmission mechanism is described using the extensively studied model introduced in [20]. In this model the behavior of uninfected, infected cells and HIV virus is given by

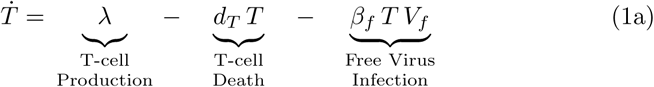

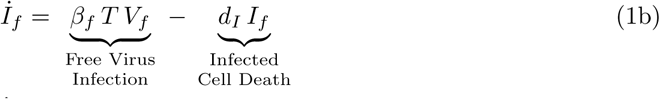

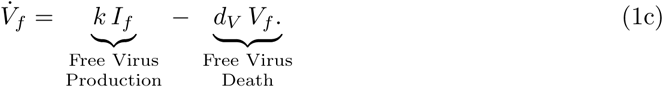

Here *T* (*t*), *I_f_* (*t*) and *V_f_* (*t*) represent uninfected cells, infected cells and free virus populations. The rate of production of uninfected cells is represented by λ. Death rates of uninfected cells, infected cells and free virus are *d_T_*, *d_I_* and *d_V_* respectively. *k* represents the number of free virus particles produced per infected cell per time unit. The mass-action infection rate is given by *β_f_*. Table 1 shows these parameters and nominal experimentally-fitted values for them obtained in [16].

If we assume that half-life of free virus is much smaller than that of infected cells ( *d_V_* >> *d_I_*), then the virus population can be assumed to be approximately in quasi-steady state, and 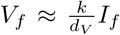. Thus Equation (1) reduces to

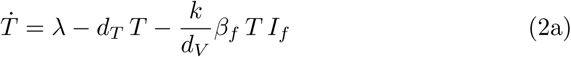

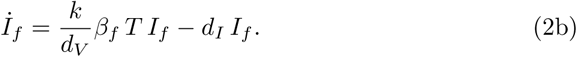

**Table 1:** All parameters values except *β_s_* were taken from [16]. *β_s_* was estimated by assuming the steady-state rate of infections by the free virus pathway is 20 times greater than rate of infections by the synaptic pathway.

| Parameter | Value | Units | Biological meaning |
| --- | --- | --- | --- |
| $\lambda$ | $7 \times 10^2$ | $\frac{cells}{\mu L \times day}$ | Uninfected birth rate |
| $d_T$ | 0.1 | $\frac{1}{day}$ | Uninfected death rate |
| $d_I$ | 1 | $\frac{1}{day}$ | Infected cells death rate |
| $d_V$ | 23 | $\frac{1}{day}$ | Virus death rate |
| $k$ | $2 \times 10^3$ | $\frac{copies}{cell \times day}$ | Copies of virus per cell |
| $\beta_f$ | $2 \times 10^{-6}$ | $\frac{mL}{copies \times day}$ | Rate of uninfected-virus interaction |
| $\beta_s$ | $10^{-5}$ | $\frac{\mu L}{cell \times day}$ | Rate of infected-uninfected interaction |

The basic reproductive ratio of infection by the free virus pathway is given by:

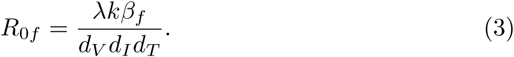

If *R*_0_ *>* 1 then the disease free condition of Equation 2 is unstable and the infection will converge to an infected steady-state.

## Modeling Synaptic Virus

Equation (1) describes transmission by free pathway. However that is not the only method of HIV transmission. Infection may also occur through direct interaction between cells, a process called synaptic transmission. When T cells come into close contact with other T cells, they sometimes form structures known as viral synapses [9]. When infected and uninfected cells form synapses, this facilitates the transfer of a large number of virions from the former to the latter [8, 9].

We modify Equation (1) in order to model synaptic transmission. Let *V_s_* (*t*) and *I_s_* (*t*) be population of free virus originating from a cell population capable of forming synapses and infected cells capable of forming a synapse at time *t*, respectively. Also let *s* be the synaptic size, which is the number of virions transmitted through a single synapse. Infected cells can now be formed through free virus infection at a rate *β_f_ TV_s_*, as well as through synapse formation at a rate *p* (*s*) *β_s_TI_s_*. Here *β_s_* is the rate of interaction between infected and uninfected cells. The function *p* (*s*) is the probability that an uninfected cell will become infected after receiving *s* virus particles through a synapse. The probability *p* (*s*) is defined as

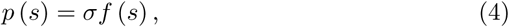
where *σ* is the probability that uninfected and infected cell form synapses given there is interaction, *f* (*s*) is the probability that sending *s* viruses through given synapses leads to an infection, and can be any monotonically increasing function on *s*. If we assume that each of *s* copies has an independent chance of successfully infecting the host cell, with each virus copy having probability *r* of successful infection then *f* (*s*) has the form:

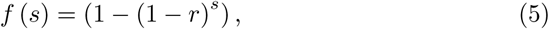
i.e. *f* (*s*) is the probability that at least one of *s* virus has successful infection given there is synapses.

There are two possible scenarios for synapse formation: infected cells with uninfected cells and infected cells with infected cells. The former leads to an infection with probability *p* (*s*). Therefore there is a reduction of *sσβ_s_T I_s_* virus copies that cannot be used in further infections. The other scenario arises because there is no known discrimination mechanism that leads infected cells to form synapses with uninfected cells only, thus infected-infected interactions also should occur. Infected-infected synapses lead to a waste of 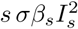 virus copies that does not produce any additional infection, because both cells are already infected. Figure 1 shows all three possible infection pathways: free virus transmission, infected-uninfected and infected-infected virus transmission through synapses.

**Figure 1:**
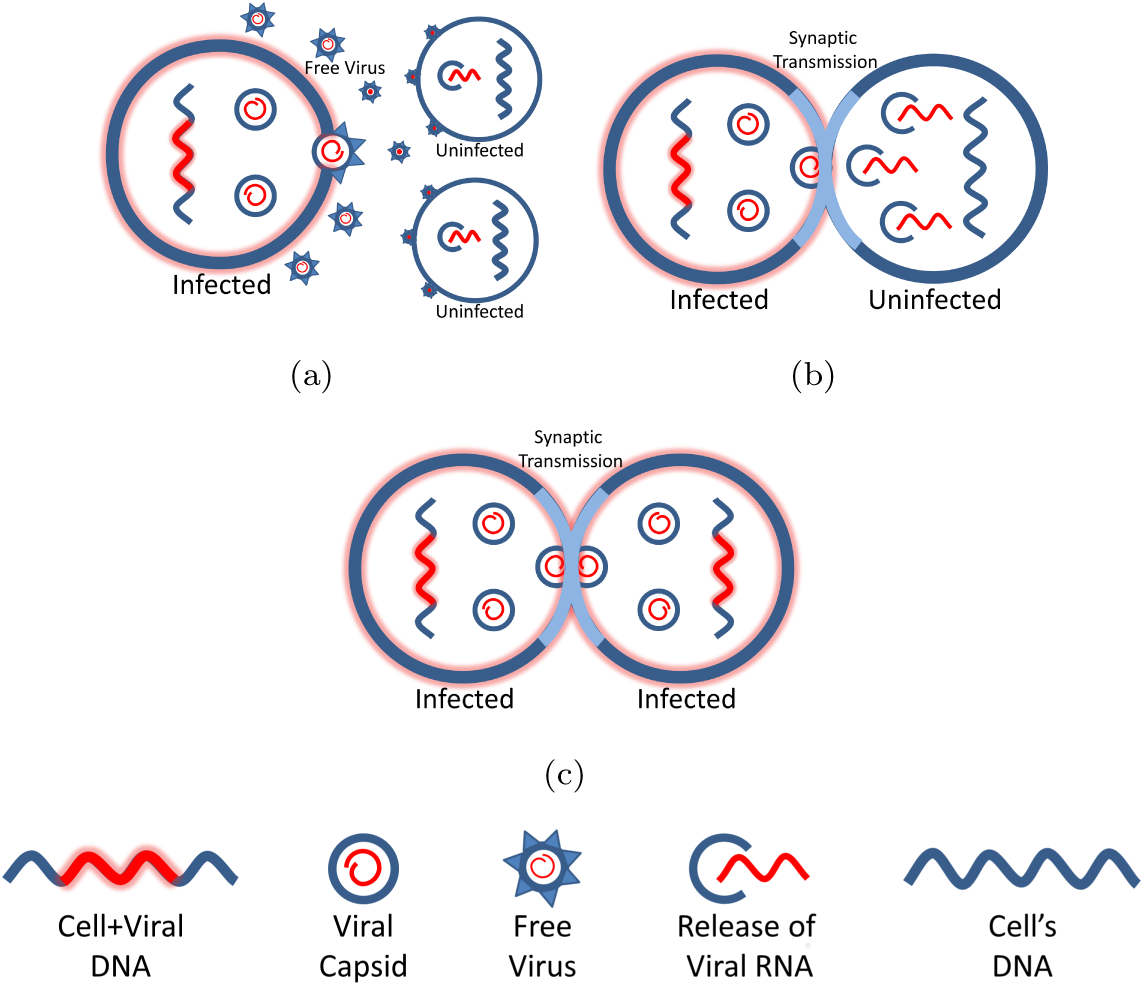
Synaptic virus mechanism. A synaptic virus has the capability of infect cells by means of free pathway (a) and also through synapses formation (b). In the free pathway (a), infected cells produce RNAs (red lines) using virus information stored in its genome (blue and red line), encapsules them (blue and red concentric circles) and send this capsids outside the cell. Uninfected cells absorb them releasing virus RNAs (opened blue circle) which integrates with cell’s DNA (blue line). Synaptic interactions may occur between infected and uninfected (b) or infected-infected cells (c). The virus copies in (b) sent through synapses are not used in the infection of other cells.

Using the synaptic mechanism illustrated above and including it in (1) leads to

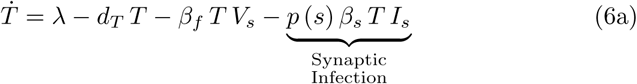

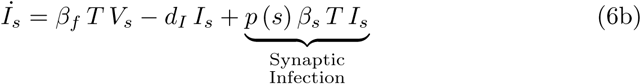

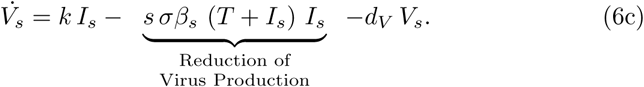

Again, if we take advantage of the fact that free virus copies die at a much greater rate than infected cells (*d_V_* >> *d_I_*), then the virus is almost always in quasi-steady state 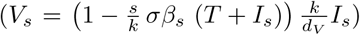 and Equation (6) reduces to

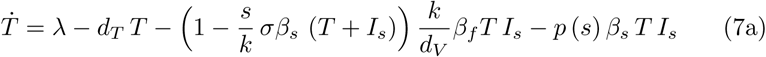

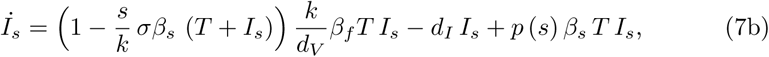
which have two stationary points, one of them being the uninfected state

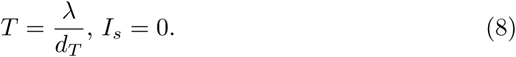

Infection will occur (this point is unstable) if

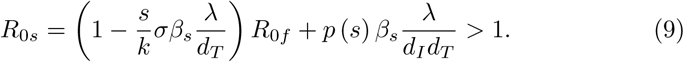

The other stability point is not difficult to calculate, however is not included here due space limits.

## Modeling effects of synaptic transmission mechanisms on HIV Latency Reservoirs

Single infection produced by the free pathway leads to two distinct situations for the new infected cell: transforms it into an active cell or latent infected cell. In the active state, the infected cell produces new viruses until it dies (lysis). When it goes latent, this infected cell does not produce new viruses. Adding the synaptic transmission mechanism can dramatically reduce the production of latent cells. This is because it allows multiple infections in a single cell, which proportionally increases the bolus of the HIV molecule Tat which is transmitted. According to the hypothesis that the cell-fate decision of a newly infected cell to become active or latent is largely due to the stochastic presence or absence of sufficient Tat to promote the early transcription of HIV, this should decrease the probability of a cell infected by the synaptic pathway of becoming latent. Figure 2 describes the possible outcomes of infection for both the free virus and synaptic transmission mechanisms.

**Figure 2:**
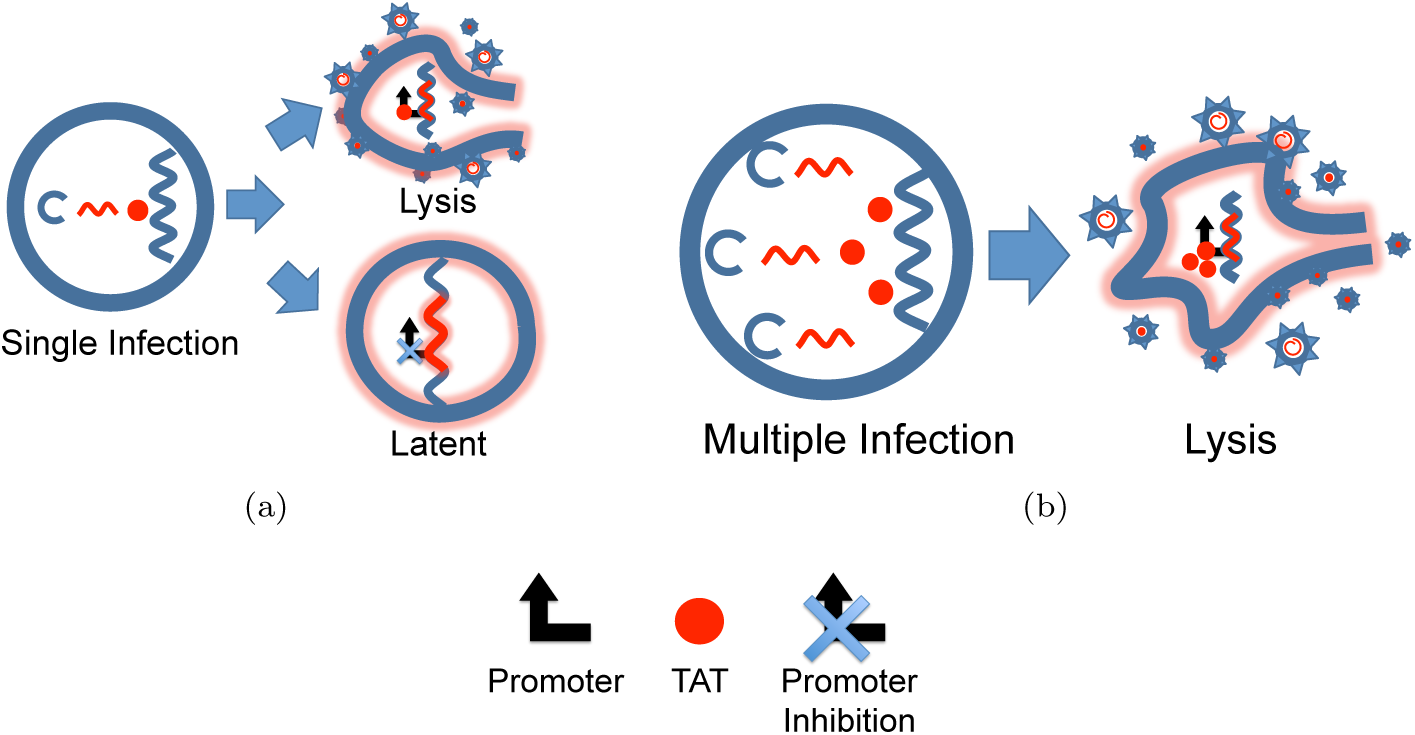
Effects of free and synaptic mechanisms on the latent pool. Free virus transmission mechanism results in a single infection with low Tat copy number that may lead the production of new viruses and subsequent lysis of the infected cell or the formation of a latent infection (a). Synaptic transmission results in a multiple infection with high Tat copy number that reduces the probability of producing latent infection (c).

In order to study how the synaptic mechanism affects HIV persistence, we add a latent infected cells pool *L_s_*. Synaptic infection effect splits now into two pools: the active, with probability

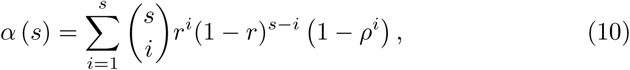
and the latent, with probability

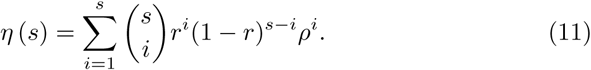
*α*(*s*) and *η*(*s*) are the probability of s or fewer virus particles sent through synapses produce an active and a latent infected cell, respectively, *ρ* is the probability that a cell infected with a single viral copy will become latent. The probabilities take this form based on the assumption that all of the independent infection events occurring with probability *r* would have a probability *ρ* of producing a latent infection if the infection is successful, but if any of them do not produce a latent infection, the outcome is an actively infected cell. Note that *α*(*s*) *+ η*(*s*) *= f*(*s*). The latently infected cells are also targets for formation of dead-end synapses that waste virus as mentioned before; it is assumed that infection of these cells does not affect their state of latency. Including all these new assumptions into equation (6) leads to the system

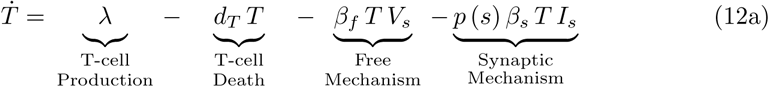

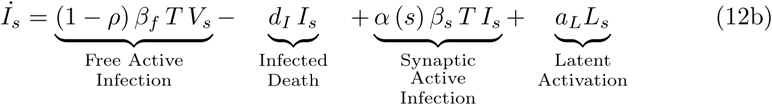

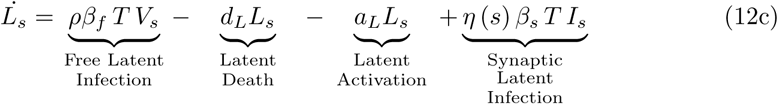

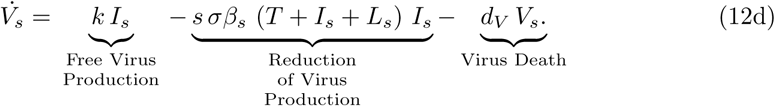

## Evaluating Fitness

The above equations describing the regulation of latency via synaptic virus transmission are analyzed to determine the conditions where synaptic transmission gives a fitness benefit to the virus. The standard measure for fitness, the basic reproduction ratio *R*_0_, is not appropriate in this circumstance, as the mechanism by which latent virus gains an advantage depends on the behavior as target cells become scarce, and *R*_0_ considers only the condition when target cells are at maximum abundance. As shown in Figure 3, it is quite easy to show that for many values of the parameters the virus with the synaptic mechanism has a lower *R*_0_, and yet outcompetes the virus without the synaptic transmission mechanism, and eventually drives it extinct.

**Figure 3:**
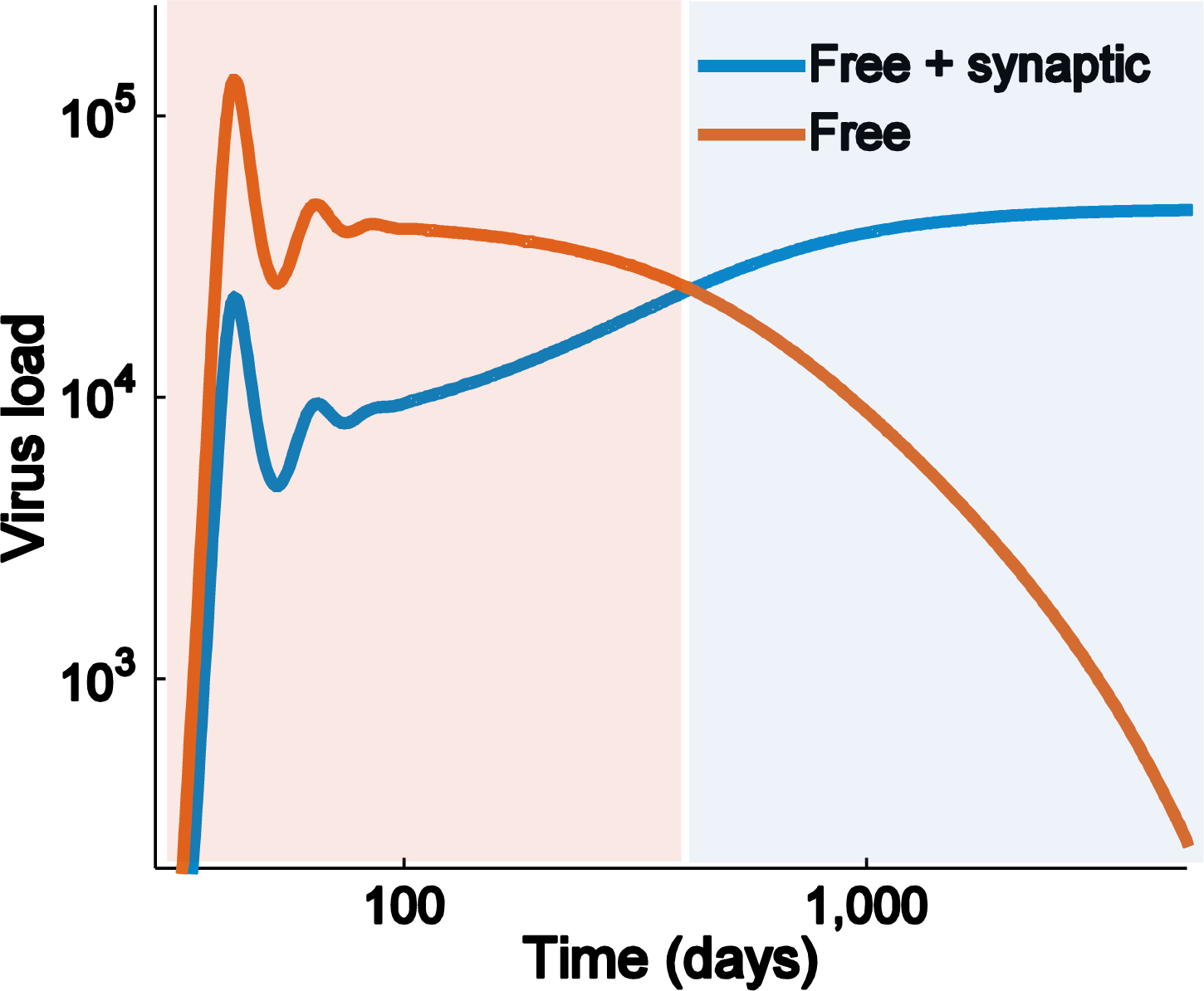
Although *R*_0 *f*_ > *R*_0 *s*_ (2 > 1:88), the synaptic virus invades the free virus. *R*_0_*f*, *R*_0_*s* are obtained using equations (5) and (11), respectively. In this case, viral fitness is determined by steady-state virus load instead of *R*_0_.

For these reasons, instead of *R*_0_, we use the steady-state infected cell count as our measure of fitness. Using this fitness criterion, the ability of the virus to devote some of its virus production to the synaptic transmission pathway shows a clear evolutionary benefit, which is stronger when the *R*_0_ of the free virus pathway is smaller. Figure 4 shows the ratio of steady-state virus loads for free virus *R*_0_ values of 1.2, 4, and 10 plotted against the fraction of virus produced which are used in synaptic transmissions.

The fitness benefit reaches a maximum for moderate levels of synaptic transmission, representing between 1% and 10% of the total virus produced; the exact maximum synaptic fraction and the actual benefit obtained depends primarily on the free virus *R*_0_ and the probability *ρ* that a cell infected by a single virus enters latency. The greatest benefit is seen for small free virus *R*_0_ and high *ρ*, whereas for large free virus *R*_0_ and high *ρ* no benefit is obtained via the synaptic virus mechanism. The dependence on *ρ* illustrates the benefit obtained by synaptic transmission through its ability to modulate the percentage of infected cells entering latency, and reducing this fraction when target cells are abundant.

**Figure 4:**
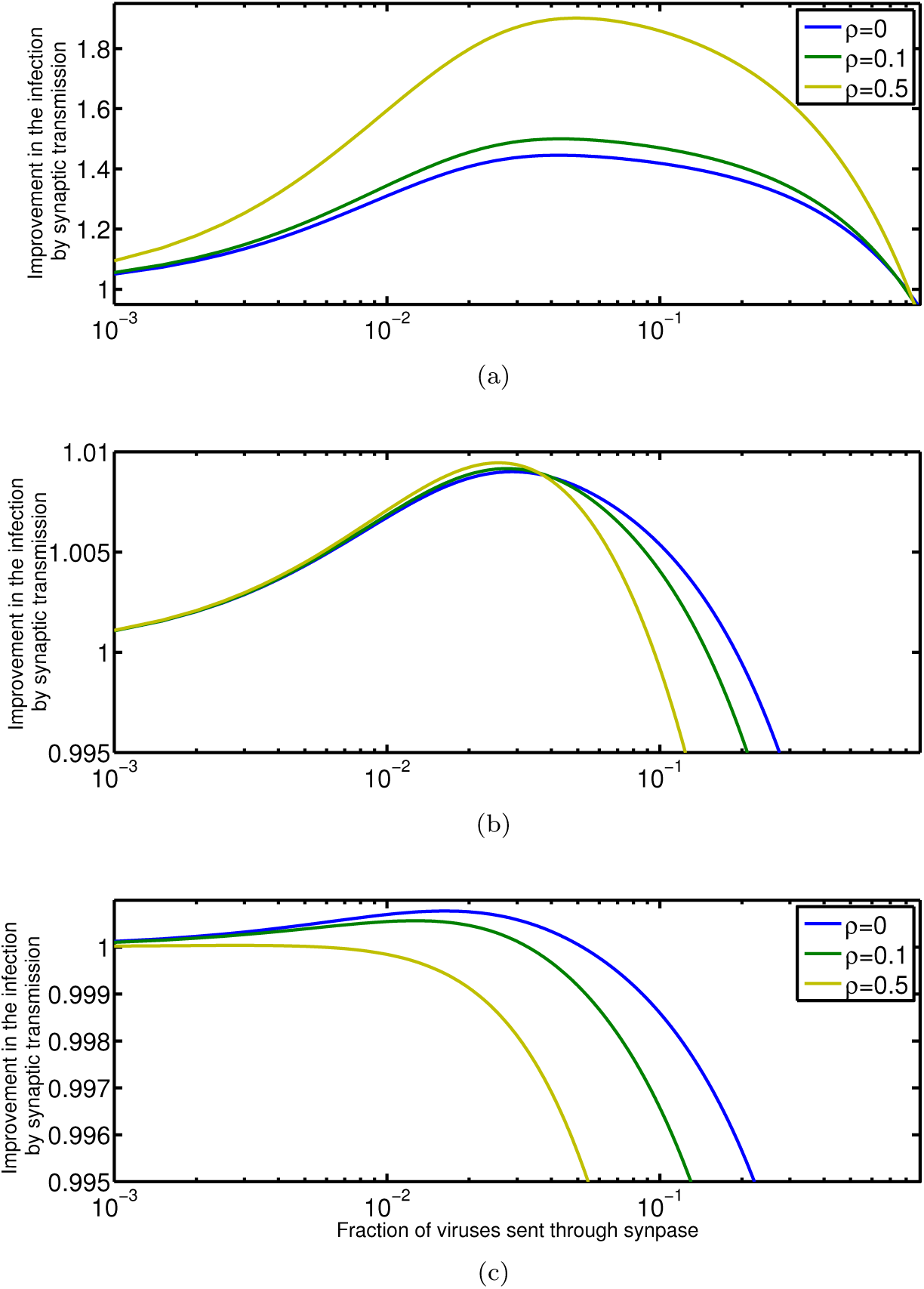
The ratio of infected cell count at equilibrium for a virus capable of synaptic transmission to a virus incapable of synaptic transmission as a function of the fraction of total virus production committed to synaptic transmission, where the basic reproductive ratio of the synaptic transmission-free virus is (a) 1.2, (b) 4, (c) 10. The value of r is 0.05 in all cases.

We also analyze the invasion criterion of the synaptic virus: the parameter values for which synaptic transmission provides an evolutionary benefit and would therefore invade a population of virus incapable of forming synapses. Analysis shows that the invasion criterion is independent of the per virion infection probability *r*, but depends on the probability of latency for a single virus infection *ρ*, the fraction of viruses sent via the synaptic pathway, and the infectivity ratio of the free virus pathway *R*_0_. These threshold values are illustrated in Figure 5 (a). For any values of the fraction of virus sent through the synaptic pathway below these critical values, the synaptic pathway provides an evolutionary benefit over the free virus pathway alone. This shows that an evolutionary benefit persists even when a remarkably high fraction of the virus production is dedicated to the synaptic transmissions. The optimal value for the fraction of virus dedicated to synaptic transmission is much lower, however, and depends greatly on *r*; this is shown in Figure 5 (b) for several values of r and a fixed value of *ρ* = 0.01.

**Figure 5:**
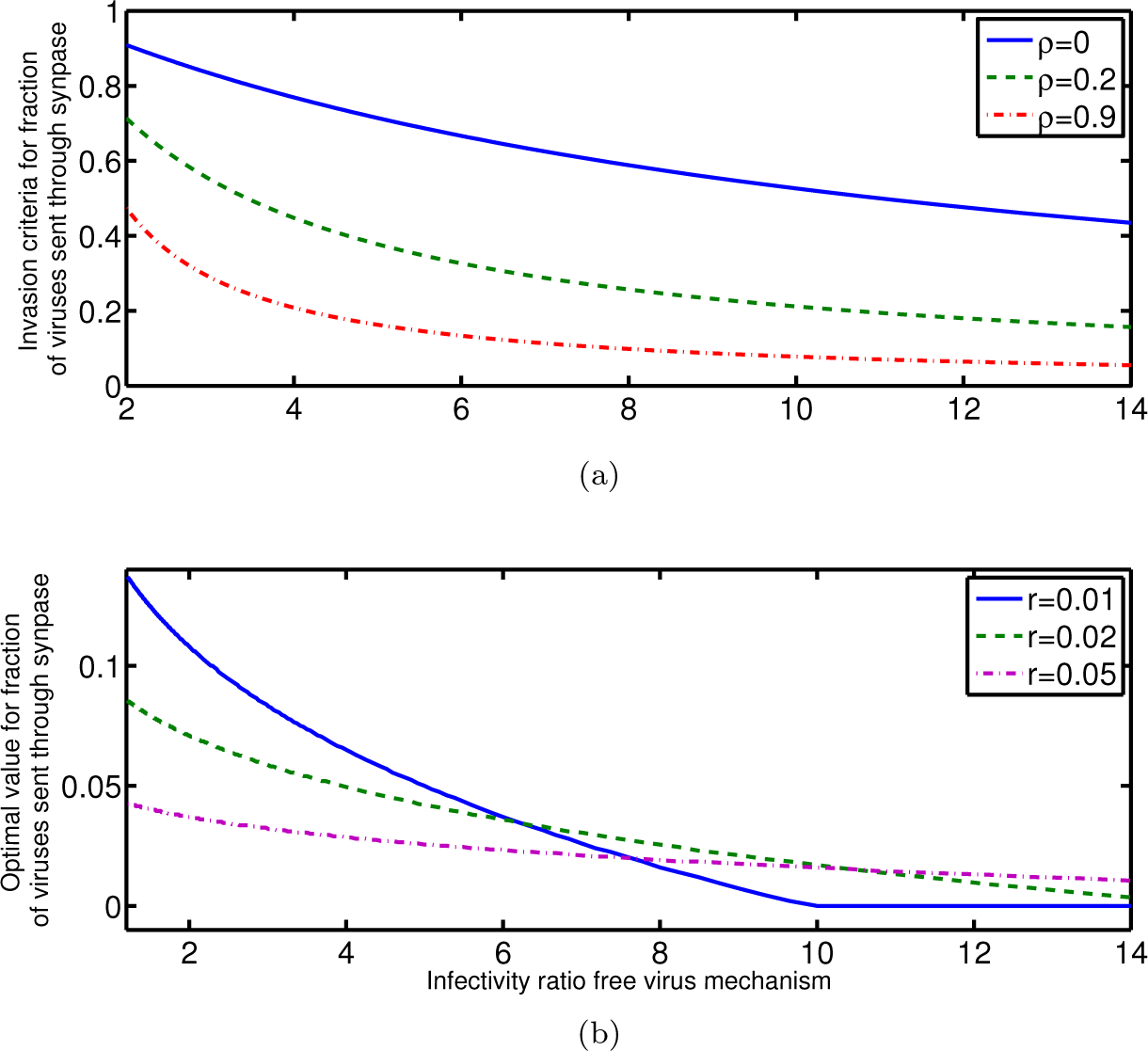
The invasion criterion (a) and optimal value (b) for the fraction of total virus production committed to synaptic transmission is plotted versus the infectivity ratio of the free virus mechanism. The invasion criterion is independent of *r*, and is plotted for different values of *p*. For panel (b), *p*=0.01.

## Conclusions

We have introduced a novel mathematical model of synaptic virus transmission and analyzed its behavior across a range of possible parameter values. This model shows that synaptic transmission provides an evolutionary benefit despite decreasing the basic reproductive ratio. This benefit increases with decreasing free virus reproductive ratio *R*_0_ and has a complex relationship with the probability of latency *ρ* and the per virion infection rate *r*, with the sensitivity of the evolutionary benefit to these parameters changing sign as *R*_0_ increases. Of particular interest is the consistent trend shown in Figure 5 where decreasing *R*_0_ results in increases both in the optimal fraction of virus committed to the synaptic pathway and the evolutionary benefit of the synaptic pathway. This may explain an observed feature of within-host evolution of HIV. During early HIV infection, the measured *R*_0_ of the virus is very high, with estimates ranging between 8 and 20 [22], which reduces to between 2 and 3 during chronic infection [16]. Furthermore, viral phenotype during transmission and early infection is nearly always dominated by a CCR5-tropic phenotype [17], but untreated infections follow a predictable pattern of within-host evolution, with the virus becoming dominated by strains that use the CXCR4 co-receptor [26]. CXCR4-tropic virus strains are associated with increased formation of tight junctions and syncytia [32], which facilitate cell-cell transmission [31];they are also associated with more rapid declines in CD4+ T-Cell counts and overall disease progression [2]. While both CCR5-tropic and CXCR4-tropic virus can both facilitate cell-cell transmission to a degree, we hypothesize that the strong preference for CCR5-tropic during transmission and acute infection may be at least partially attributed to the evolutionary disadvantage associated with the cell-cell pathway when the free-virus *R*_0_ is high, and that the subsequent pattern of evolution of co-receptor diversity may be attributed to the shift toward a strong evolutionary benefit for the cell-cell pathway once the free virus *R*_0_ decreases during chronic infection.

## Acknowledgements

Research reported in the publication was supported by the National Institutes of Health (NIH) grant AI110288. The content is solely the responsibility of the authors and does not necessarily represent the official view of the funders. The funders had no role in study design, data collection and analysis, decision to publish, or preparation of the manuscript.

